# Characterization of the *Tuta absoluta* virome reveals higher viral diversity in field populations

**DOI:** 10.1101/2024.12.16.628664

**Authors:** Rosa Esmeralda Becerra-García, Luis Hernández-Pelegrín, Cristina Crava, Salvador Herrero

## Abstract

A significant number of insect-specific viruses (ISVs) have been discovered in agriculturally important insect pests, facilitated by high-throughput sequencing (HTS). Despite its global impact on tomato crops, the RNA virome of the South American tomato pinworm, *Tuta absoluta*, remains uncharacterized. In this study, we utilized meta-transcriptomics and bioinformatic approaches to discover the RNA virome of *T. absoluta* across worldwide populations. We identified ten novel ISVs, classified into six groups: Nidovirales, Bunyavirales, Mononegavirales, *Virgaviridae*, *Iflaviridae*, *Nodaviridae*, *Solemoviridae*, and *Phasmaviridae*. Notably, no core virus was consistently present across the studied populations, and field-collected samples revealed a greater diversity of ISVs compared to those from laboratory samples. In addition, we detected plant-infecting viruses and mycoviruses associated with the pest. This study represents the first description of the RNA virome associated with *T. absoluta*, providing valuable insights into its biological and ecological interactions. It also lay the foundation for future studies aimed to clarify the biological roles of ISVs.

## Introduction

The tomato leafminer or tomato pinworm *Tuta absoluta* (Meyrick) (Lepidoptera: Gelechiidae), is a microlepidopteran considered one of the most destructive pests affecting tomato crops worldwide. Originally from Peru, *T. absoluta* entered Europe in 2006 and has rapidly spread to over 90 countries outside from South America (Desneux *et al*., 2022). Several factors classify *T. absoluta* as a significant pest, including its relative short life cycle, which allows for multiple generations within a year, its behavior of mining and feeding on the mesophyll of tomato plants, creating galleries that can lead to the destruction of entire plants, its rapid spread, and its resistance to insecticides (Desneux *et al*., 2022). This pest also attacks other Solanaceous plants, such as potato (*Solanum tuberosum*) and black nightshade (*Solanum nigrum*) (Abbes *et al*., 2016).

Since the 1960s, *T. absoluta* has been primarily controlled using chemical insecticides. Throughout the years, however, the increasing resistance to these insecticides has necessitated the development and adoption of alternative control methods. Nowadays, some of the most effective management practices behind chemical insecticides include trapping, the use of sex pheromones, and biological control strategies involving both microorganisms and arthropods (Guedes *et al*., 2019, Desneux *et al*., 2022). Among the microorganisms used as biological control agents is the Phothorimaea operculella granulovirus (PhopGV), a DNA virus isolated from various populations of the potato tuber moth *Phothorimaea operculella* worldwide. PhopGV is pathogenic to *T. absoluta* and cause significant mortality in its larvae (Gómez Valderrama *et al*., 2018, Ben Tiba *et al*., 2019). However, no information is currently available on other viral species infecting this insect or their potential implications for its ecology and biological control.

Over the past decade, the application of metagenomic and metatranscriptomics approaches has led to the discovery of numerous novel RNA Insect-Specific Viruses (ISVs), which are unique in that they exclusively replicate within insect cells (Li *et al*., 2015, Shi *et al*., 2016, Käfer *et al*., 2019, Wu *et al* 2020, Qi *et al*., 2023). Studies have revealed that the most abundant viral families in agricultural pests include *Iflaviridae*, *Dicistroviridae*, *Partitiviridae*, and *Rhabdoviridae* (Wu *et al* 2020, Qi *et al*., 2023). In the last years, these RNA viruses have gained increasing attention due to their ability to cause both covert and overt infections in their hosts (Liu, *et al*., 2011, Ryabov, 2017). This duality positions them as potential tools for biological control (Nouri *et al*., 2018, Bonning, 2019). For instance, within lepidopteran insects, iflaviruses exhibit a range of effects, from pathogenic (Geng *et al*., 2014; Silva *et al*., 2015) to non-pathogenic (Millán-Leiva *et al*., 2012; Choi *et al*., 2012). In *Spodoptera exigua*, the iflavirus 1 (SeIV1) is not lethal but significantly increases the susceptibility of the insect to the Spodoptera exigua multiple nucleopolyhedrovirus (SeMNPV), a baculovirus widely used in biological control (Jakubowska *et al*., 2016; Carballo *et al*., 2017, 2020). This virus also increases the insect’s vulnerability to the entomopathogenic bacterium *Bacillus thuringiensis* (Mengual-Martí *et al*., 2022). Other RNA ISVs, such as partiti-like viruses, also play notable roles in host dynamics (Xu *et al*., 2020). Three of these viruses, identified in the African armyworm (*S. exempta*) have been shown to reduce the growth rate and reproduction of the host while simultaneously increasing its resistance to a nucleopolyhedrovirus (Xu *et al*., 2020). These findings emphasize the complex interplay between ISVs and their hosts, highlighting their potential implications for pest management strategies.

In this context, our study provides the first comprehensive characterization of the RNA virome of *Tuta absoluta*, based on metatranscriptomics and bioinformatic analyses of both newly generated transcriptomes and publicly available Sequence Read Archive (SRA) datasets. We identified ten ISVs, along with plant-infecting viruses and mycoviruses associated with this insect, revealing a large heterogeneity in the viral profile across different global populations.

## Materials and methods

### Field collection of Tuta absoluta

Larvae and moths of *T. absoluta* were collected from infested *Solanum lycopersicum* tomato plants in different locations of Spain from both greenhouses and open field crops. Leaves of *S. lycopersicum* that were infested with *T. absoluta* were detached from the plant and live or dead larvae were manually sorted and stored at −80°C until further processing. Larval samples were collected from Almeria, Xilxes, Blanes and Quintana, and Zafarraya and analyzed independently. In contrast, adults were sampled in Agua Dulce, Níjar, Vicar, Campo hermoso, and Alquian and analyzed as a pool named Multiple locations (ML). Additionally, larvae from a laboratory colony, originally established from insects collected in greenhouses of Peñiscola (Spain) in 2015 and reared at the IVIA (Instituto Valenciano de Investigaciones Agrarias), were included as an independent sample (Suppl Table S1).

### RNA extraction and sequencing

In total, individuals from different locations and grouped according to their health status were initially pooled. Each pool contained between five to thirty larvae or moths. Frozen pools were homogenized in 200 µl PBS 1x with a pestle and 50 µl of the homogenates were used for total RNA extraction using 500 µl TRIzol reagent and following the manufacturer’s instructions. RNA was quantified using a NanoDrop Spectrophotometer (TermoFisher) and the integrity was visualized in a 1% agarose gel. Pools were sent to Macrogen, Inc (Republic of South Korea) for DNAse treatment, ribosomal RNA depletion, cDNA library preparation, and high-throughput sequencing. Illumina NovaSeq 6000 paired-end sequencing was used to generate 150bp paired-end RNA-Seq reads. Raw data are available in NCBI SRA (Sequencing Read Archive) database under the BioProject accession number PRJNA1188764.

### Transcriptome Assembly and Virus Discovery

For the virus discovery, a pipeline previously described (Hernández-Pelegrín *et al*., 2022) was followed, which consists in three main steps: quality-control of the paired-end reads, assembly of the reads and detection and annotation of viral sequences.

The quality of the paired-end RNA-seq reads were analyzed using FastQC (https://www.bioinformatics.babraham.ac.uk/projects/fastqc/, accessed on April 6^th^ 2023), adaptors were removed, and low-quality sequences were trimmed using Trimmomatic (Bolger *et al*., 2014). The RNA-seq reads of 16 meta-transcriptomic pools of *T. absoluta* were *de novo* assembled using Trinity-v2.15.1 (Grabherr *et al*., 2011). All programs were used with default parameters. To detect viral sequences in the assembled contigs, the tBLASTn (protein query against translated nucleotide database) analysis was used and restricted with an e-value threshold of 1 x 10^−5^. As the query, protein viral sequences deposited in GenBank were employed, categorized within the Riboviria domain. Following tBLASTn analysis, putative viral contigs were manually filtered, excluding those based on three criteria: contigs less than 1000 bp in length, contigs identified in the reference genome of *T. absoluta* after BLASTn (e-value cut off of 1×10^−100^), and contigs without open reading frames (ORFs) after evaluation with ORFfinder considering a minimal ORF length of 150 nucleotides and alternative initiation codons (https://www.ncbi.nlm.nih.gov/orffinder/, accessed on May 5^th^, 2023).

To determine the presence of viral sequences in transcriptomes of worldwide *T. absoluta* samples, we used the Serratus platform (https://serratus.io/explorer/rdrp, Edgar *et al*., 2022) to evaluate 42 SRA datasets from six bioprojects encompassing populations from Europe, South America and Asia, available in NCBI database (Suppl. Table S1). Those SRA files with hypothetical RdRp domain were further assembled and analysed for viral discovery as described above.

### Annotation of Viral Sequences

Putative viral sequences were annotated by conducting BLASTx analysis against the non-redundant (nr) protein sequence database of NCBI. In addition, the ORFs of the viral sequences were identified, followed by BLASTp analysis against the nr protein sequence database and restricted to “viruses” to identify conserved domains for each ORF. The ORFs were also evaluated using the InterPro tool (https://www.ebi.ac.uk/interpro/) (Paysan-Lafosse *et al*., 2023). Viral names were assigned as: name of the host, followed by the root of the viral family, plus virus, and an identification number ordered by nucleotide sequence length (from longest to shortest). When family demarcation was unclear, the term “like-virus” was used. Accession numbers for the nucleotide sequences of the new viruses described in *T. absoluta* are included in Suppl. Table S2.

### Phylogenetic Analysis

Phylogenetic analyses were conducted using the protein sequences of conserved domains, mainly based on RNA-dependent RNA polymerase (RdRp). In some cases, the Helicase (Hel) or Coat protein (CP), depending on available information of the viruses under study, was used. The closest viral relatives for the identified viruses were determined using BLASTp analysis against the nr database. Viruses described in lepidopteran species within the same viral families in addition to representative viruses classified by the International Committee on Taxonomy of Viruses (ICTV) were added into the analysis to support the phylogenetic inference (Suppl. Table S2). Multiple sequence alignment (MSA) was performed using T-Coffee (di Tommaso *et al*., 2011) and poorly aligned nucleotides were manually trimmed using Bioedit (Hall, 1999). The MSA was employed to infer maximum likelihood phylogenetic trees using IQtree2 and 1000 ultrafast bootstrap replicates with default parameters (Minh *et al*., 2020). Trees were visualized and edited using iTOL (Letunic & Bork, 2021) and rooted at the midpoint.

### Abundance of viral sequences

The presence and relative abundance of the discovered viruses were evaluated *in silico* using the 16 metatranscriptomes of *T. absoluta* generated in this study and 42 SRA datasets available in NCBI database. The sequencing reads were mapped against the genomic sequences of the discovered RNA viruses of *T. absoluta* using Bowtie 2 v 2.3.5.1 (Langmead & Salzberg, 2012), and viral abundance was calculated using RSEM v 1.3.1 (Li & Dewey, 2011) with default parameters. The log relative abundance of each transcript is reported relative to the *T. absoluta* endogenous gene Elongation Factor 1-alpha (*EF1α*, Genbank: MZ357901.1). The expression data obtained from the analysis were represented using the heatmap.2 function from the gplots v 3.1.3 package of the R software (Warnes *et al*., 2009).

## Results

### Diversity of RNA viruses discovered in T. absoluta

Sixteen meta-transcriptomic datasets, generated from pools of field-collected individuals, were grouped based on their location and whether they represented live and dead larvae. Following sequences assembly, viral sequences potentially representing 27 infective viruses associated with *T. absoluta* were identified. Among the viruses, seven are presumably ISVs of *T. absoluta*, while four correspond to plant-infecting viruses, and 13 are potential mycoviruses. To further expand our analysis of the ISVs in *T. absoluta*, we analyzed transcriptome data from publicly available databases derived from samples collected in Spain, Portugal, Greece, Brazil, and China. This analysis revealed three additional ISVs, increasing the total to ten ISVs associated with *T. absoluta.* These findings represent the first discovery of RNA viruses in worldwide distributed populations of the tomato leafminer (Fig. 1).

**Fig. 1.**
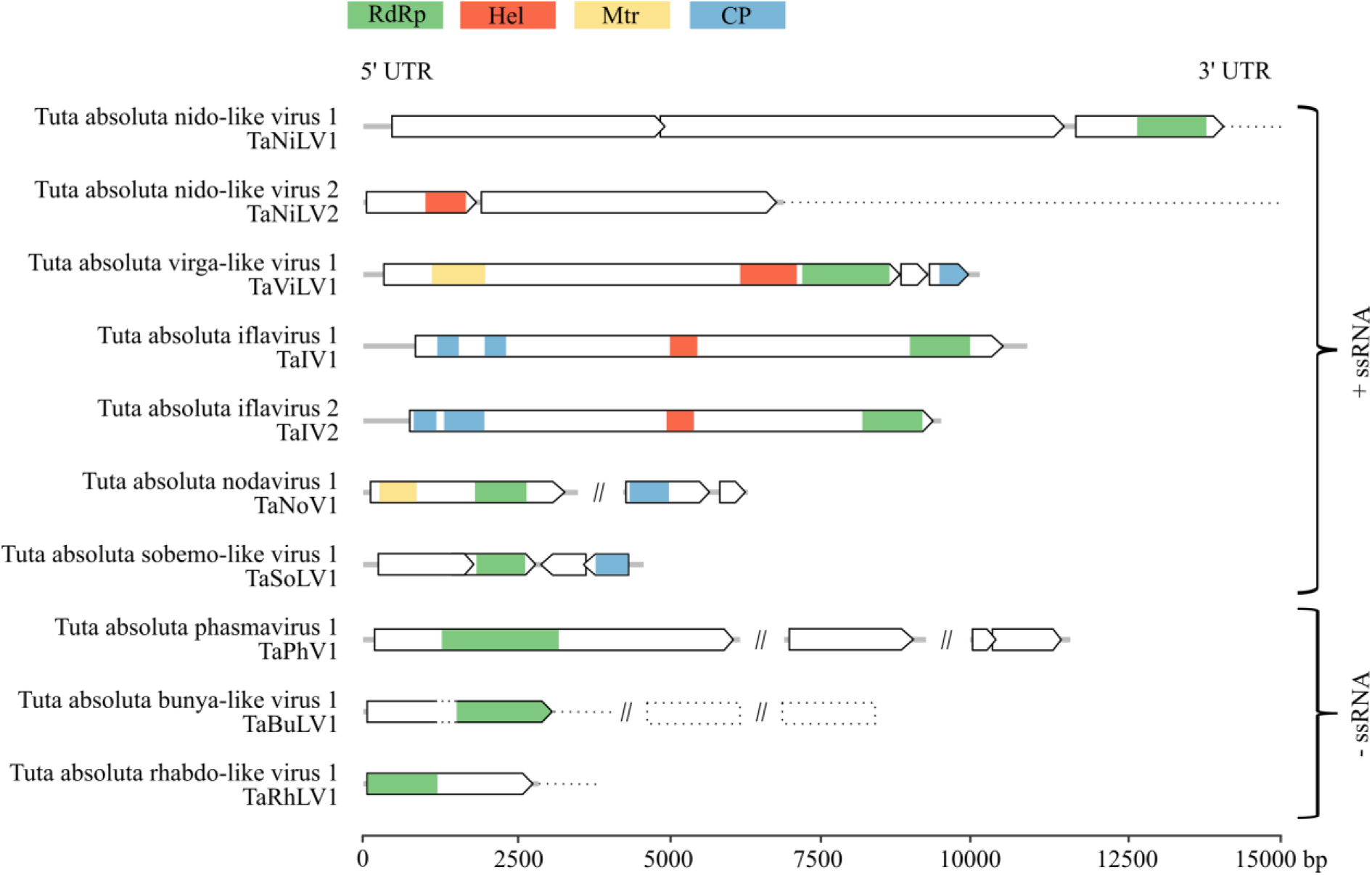
Putative genomic organization of the ten novel insect-specific viruses discovered in *T. absoluta*. White boxes represent putative ORFs, and the conserved protein domains RdRp (RNA-dependent RNA polymerase), Hel (Helicase), Mtr (Methyltransferase) and CP (Coat protein) are color-coded. Each genome is illustrated in proportion to its length. Dotted line indicates the hypothetical part of the genome that is currently missing. TaNoV1, TaPhV1, and TaBuLV1 presented a segmented genome, and double slashes separate the corresponding fragments. Six ISVs belong to positive sense single-stranded RNA (+ssRNA), whereas four belong to negative sense single-stranded RNA (-ssRNA).

Viruses were classified based on their genome structure and phylogeny. ISVs were taxonomically classified into eight groups of ssRNA viruses. Six were classified within the families *Virgaviridae*, *Iflaviridae* (two viruses), *Nodaviridae*, *Solemoviridae* and *Phasmaviridae* (Suppl. Fig. S1). Four additional viruses were classified at the order level: two belonging to Nidovirales, one to Bunyavirales, and one to Mononegavirales (Suppl. Fig. S1). The name and abbreviations of these viruses are proposed as Tuta absoluta virga-like virus 1 (TaViLV1), Tuta absoluta iflavirus 1 (TaIV1), Tuta absoluta iflavirus 2 (TaIV2), Tuta absoluta nodavirus 1 (TaNoV1), Tuta absoluta sobemo-like virus 1 (TaSoLV1), Tuta absoluta phasmavirus 1 (TaPhV1), Tuta absoluta nido-like virus 1 (TaNiLV1), Tuta absoluta nido-like virus 2 (TaNiLV2), Tuta absoluta bunya-like virus 1 (TaBuLV1), and Tuta absoluta rhabdo-like virus 1 (TaRhLV1).

Among the viruses with a +ssRNA genome, TaNiLV1 has a putative nearly complete genome of 14,060 bp, featuring three ORFs, two of which are partially overlapping. The third ORF encodes for the RNA-dependent RNA polymerase (RdRp) (Fig. 1). In contrast, TaNiLV2 has a putative partial genome of 7,398 bp, comprising two putative ORFs. The first ORF contains the helicase (Hel) protein domain, while no RdRp domain was found (Fig. 1), likely due to its absence in the sequenced region. Phylogenetic analysis of TaNiLV1 and TaNiLV2, based on RdRp and Hel proteins respectively, place them within the insect nidovirus group (Suppl. Fig. S1A). TaViLV1 presents a complete genome of 10,170 bp containing three ORFs. The largest ORF contained three protein domains: Mtr (methyltransferase), Hel and RdRp. ORF 3 is likely to encode a CP (Fig. 1). Two complete genomes of 10,273 bp and 9,463 bp were identified for TaIV1 and TaIV2, respectively. Both feature a single ORF that encodes a polyprotein containing all the functional domains typical of iflaviruses. TaNoV1 has a segmented genome, with one segment of 3,226 bp including both the Mtr and RdRp domains, and a second segment of 1900 bp containing the CP domain (Fig. 1). TaSoLV1 has a genome of 4,289 bp, composed of four ORF. The second ORF encodes an RdRp, while the fourth ORF encodes a CP (Fig. 1). Phylogenetic analyses based on RdRp proteins from the five +ssRNA viruses supported their classification as insect-infecting viruses (Suppl. Fig. S1B-E).

Three viruses with -ssRNA genomes were also identified, each with distinct features. TaPhV1 presented a complete segmented genome of 5,979 bp, comprising two segments of 2,411 bp and 1,554 bp, respectively. The longer segment encodes the RdRp function. TaBuLV1 presented a partial segment containing the RdRp domain. Based on the typical genome structure of the bunyaviruses, we suspect that two additional segments are missing. This is likely due to the low sequence homology with known viruses and insufficient sequencing coverage (Fig. 1). TaRhLV1 presented a partial genome of 2,744 bp, containing one ORF that includes the RdRp domain (Fig. 1). However, given the length of related rhabdoviruses, it is likely that approximately 5 kb are missing. Phylogenetic analysis supported the classification of the -ssRNA viruses as insect-infecting viruses (Suppl. Fig. 1F-H).

In addition to the ISVs of *T. absoluta,* the metatranscriptomic analysis of the field-collected samples in this study also revealed the presence of known plant-infecting viruses and mycoviruses. The identified plant-infecting viruses were potato virus Y (PVY), tomato chlorosis virus (ToCV), tomato brown rugose fruit virus (ToBRFV), and tomato mosaic virus (ToMV). Concerning the mycoviruses, the analysis led to the discovery of six new hypothetical mycoviruses, which were provisionally named until their hosts are confirmed as: Tuta absoluta-associated mitovirus 1, Tuta absoluta-associated narnavirus 1, Tuta absoluta-associated ourmiavirus 1, Tuta absoluta-associated ourmiavirus 2, Tuta absoluta-associated ourmiavirus 3, and Tuta absoluta-associated bunya-like virus 1 (Suppl. Fig S2). Phylogenetic analysis suggested that these viruses were more related to mycoviruses than to insect viruses (Suppl. Fig S1G,I). Additionally, seven other detected viruses were previously identified as associated with fungal and protist species (Suppl. Fig. S2).

### Prevalence, abundance, and distribution of the RNA viruses

After virus annotation, the prevalence and abundance of all discovered viral sequences were evaluated across the 16 transcriptomes generated in this study, which represent samples from different geographical field locations and larval health status (live vs dead larvae, Fig. 2, Suppl. Table S1). Additionally, we screened their occurrence in 42 SRA datasets available in the NCBI database, encompassing populations from Europe, South America, and Asia (Fig. 2, Suppl. Table S1). In total, all the 58 transcriptomes evaluated represented 14 different populations.

**Fig. 2.**
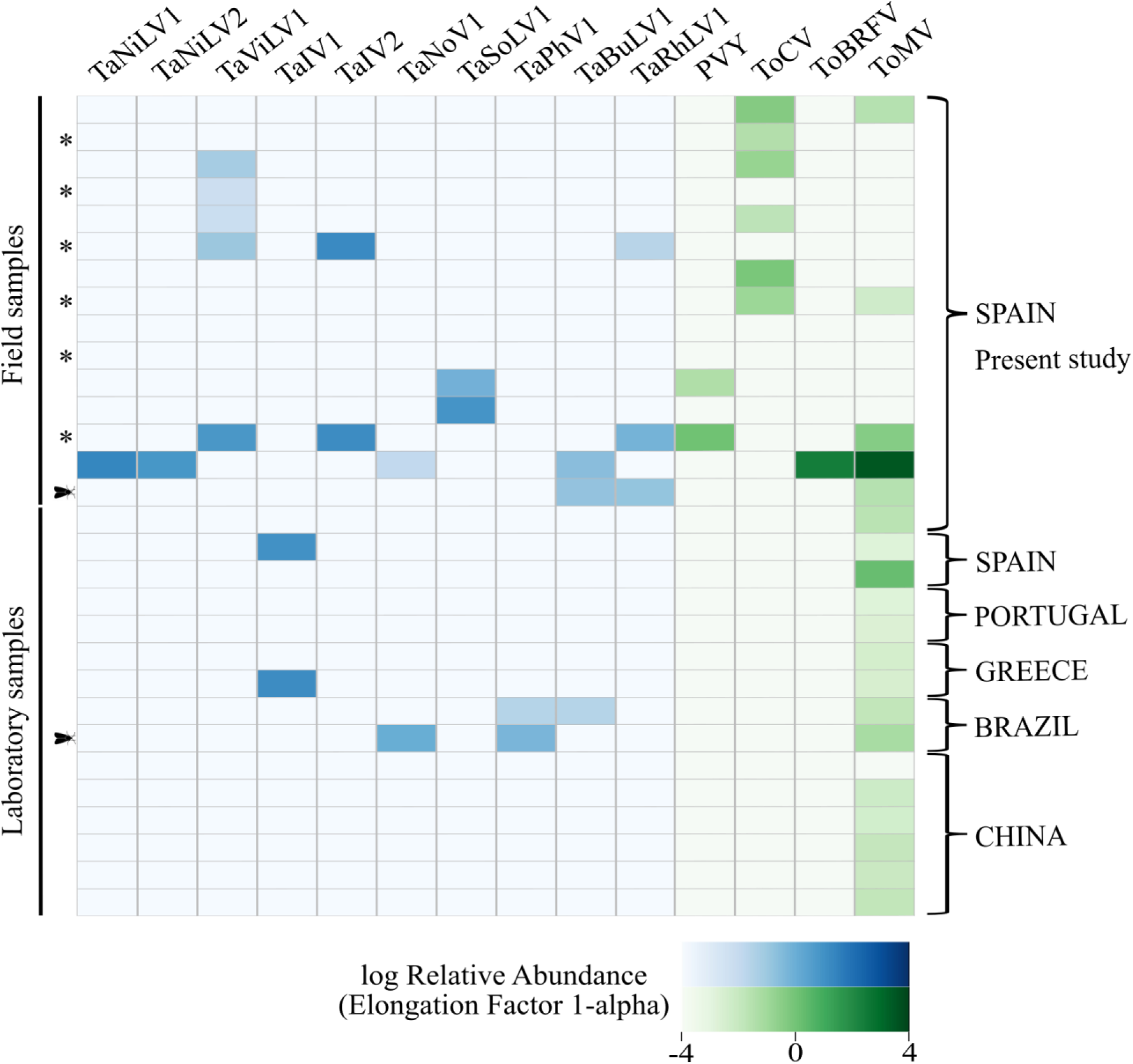
Relative abundance of discovered RNA viruses in different *T. absoluta* samples. Heatmap of the relative abundance of 14 RNA viruses identified in 16 meta-transcriptomic pools, as well as 14 representative transcriptomes available in the NCBI database corresponding to samples from different global locations (for additional information see Suppl. Table S1). The viral relative abundance was obtained by normalizing to the expression of the endogenous gene *EF1α* of *T. absoluta*. Columns without symbol refers to samples composed by live larvae, asterisk symbol refers to dead larvae, and icon of moth refers to moth samples. The ISVs and plant viruses are represented in blue and green colors, respectively.

The distribution of RNA viruses varies among *T. absoluta* samples, with no single core virus present in all the samples (Fig. 2). Notably, the majority of ISVs (six) were present only in field-derived samples, whereas two ISVs were present in both laboratory and field-collected samples and other two specifically present in laboratory samples. Also, the presence of at least one viral infection was higher in field-collected samples (nine out of 15 samples with viral infection) compared to laboratory samples (four out of 15), suggesting a greater diversity of RNA viruses in samples collected from the field (Fig. 2).

Among the ISVs identified in the field-collected samples (all of them produced during this study), the most prevalent was TaViLV1, which was present in five out of 16 samples. TaRhLV1 was found in three samples of different locations whereas TaIV2, TaSoLV1 and TaBuLV1 were present in two locations each and TaNoV1 was present only in one sample (Fig. 2). For most of the ISVs present in field-collected samples, they were equally found in dead and live larvae, with the exception of TaIV2, which was exclusively found in samples prepared from dead larvae derived from two different locations (Fig. 2). Among the ISVs identified in laboratory samples, we generally found a low prevalence, with many samples totally free of viruses (11 out of 15). The highest titers were detected for TaIV1, which infected two samples from Spain and Greece, respectively. At lower titers, TaNoV1, TaPhV1, and TaBuLV1 were detected in Brazilian laboratory samples (Fig. 2). Samples from China and Portugal did not have any trace of viral infection. It is noteworthy that two field-collected samples were entirely free of any of the 27 identified viruses (including ISVs, plant-infecting viruses, and mycoviruses) (Fig.2, Suppl Fig. S2).

Regarding plant-infecting viruses, ToMV was the most prevalent virus (present in five out of 15 field-collected samples and 14 out of 15 laboratory samples), while the highest titer was displayed by both ToMV and ToBRFV in the same sample (Fig. 2). The analysis of mycoviral sequences across the 16 metatranscriptomic datasets from this study revealed their presence in only four samples. Notably, ten viruses were detected in a single sample from field-collected dead larvae (Suppl. Fig. S2).

## Discussion

In this work we characterized the first RNA virome of the tomato leafminer, *Tuta absoluta*. The analysis of metatranscriptomic data generated from field samples collected during this study, along with public transcriptomes derived from laboratory populations, led to the discovery of ten novel insect-specific viruses that infect this crop-damaging lepidopteran species. To our knowledge, no *T. absoluta*-infecting RNA viruses have been described until now. This may be due to the cryptic phenotype of the viruses, which infect apparently healthy insects without causing overt symptoms (Nouri *et al*., 2018). In this line, advances in high-throughput sequencing have greatly benefited viral discovery, enhancing our understanding of viromes in different species that previously went undetected due to their subtle phenotypic effects (Li *et al*., 2015, Shi *et al*., 2016, Käfer *et al*., 2019, Wu *et al* 2020). Notably, we identified ISVs from well-known viral families associated with covert infections in Lepidoptera, as well as uncovered ISVs from less-characterized viral families in this order, thereby expanding our understanding of the RNA virome in Lepidoptera. Examples of these covert infections include Spodoptera exigua iflavirus 1 (SeIV1) and Spodoptera exigua iflavirus 2 (SeIV2), which are commonly found in *S. exigua* populations without causing overt signs of disease (Millán-Leiva *et al*., 2012; Choi *et al*., 2012, Virto *et al*., 2014). Similarly, within the *Spodoptera* genus, Spodoptera frugiperda rhabdovirus is frequently detected in pheromone-trapped individuals, with no observed cytopathic effects in insect cell lines (Schroeder *et al*., 2019). Likewise, a bunyavirus infecting the tea tussock moth, *Euproctis pseudoconspersa*, is widely distributed in field populations across China (Wang *et al*., 2021). In addition, a lepidopteran-infecting nodavirus has been identified in the butterfly *Pieris rape* (Liu *et al*., 2006). Together, these findings demonstrate how ISVs producing covert infections can persist in natural populations without inducing major symptoms. Other viral families identified in this study, such as *phasmaviridae*, *solemoviridae*, and *nidoviridae*, are better known for their associations with Diptera, particularly mosquitoes, where they also produce covert infections (Ribeiro *et al*., 2020, Shi *et al*., 2020, Maia *et al*., 2024, Pradhan *et al*., 2024). However, there is limited knowledge about their ability to infect Lepidoptera. This knowledge gap likely arises from the extensive research on mosquito viromes due to their relevance to human health, compared to the relatively sparse studies on Lepidoptera viromes. We forecast that future studies will broaden the number of Lepidoptera-infecting viruses that are present as covert infection in specific species, mainly agricultural pests, since it is an essential first step toward understanding the biological roles these viruses play in insect ecology and their potential applications in biological control.

The viral profile of *T. absoluta* is heterogeneous among the samples, indicating the absence of a core ISV. Eight out of ten viruses were detected in field-collected samples from Spain, whereas only four were present in laboratory samples, highlighting a higher diversity of RNA viruses in field-collected samples. For Lepidoptera, viral screenings of *S. exempta* and *S. frugiperda* from laboratory colonies did not reveal viral infections, despite their detection in field-collected individuals (Xu *et al*., 2020). Similar scenarios have been reported for Diptera, where higher viral diversity was also observed in field-collected samples compared to laboratory samples (Shi *et al*., 2020, Hernández-Pelegrín *et al*., 2022).

Both TaNoV1 and TaBuLV1 were detected in samples from both field and laboratory environments. Remarkably, in our study, TaIV1 was found in samples from both Spain and Greece, being abundant in laboratory samples but absent in field-collected samples. Conversely, TaIV2 was exclusively found in dead larvae from field-collected samples, suggesting a potential pathogenic role in *T. absoluta*. However, further studies are required to confirm this lethal effect, which has been observed in other iflaviruses. For instance, the iflavirus infecting the Chinese oak silkmoth, *Antheraea pernyi*, causes the *A. pernyi* vomiting disease (AVD), where infected larvae exhibit sluggish behaviour and a white liquid vomited from the midgut before ceasing to feed (Geng *et al*., 2014). Likewise, the Opsiphanes invirae iflavirus 1 (OilV-1), found in dead larvae of *Opsiphanes invirae*, a common pest of the African oil palm tree, is pathogenic to its host (Silva *et al*., 2015). This highlights that while some iflaviruses may be directly lethal, others can influence host-pathogen interactions, for instance, by affecting the susceptibility of the host to entomopathogens (Jakubowska *et al*., 2016; Carballo *et al*., 2017, 2020, Mengual-Martí *et al*., 2022).

The RNA viruses TaNoV1, TaPhV1 and TaBuLV1 were identified in a *T. absoluta* population from South America, its native region (Guillemaud *et al*., 2015, Biondi *et al*., 2018). Of these, only TaPhV1 was exclusively detected in this South American population. Interestingly, most of the new RNA viruses identified in this study were unique to samples collected in Spain and were absent in laboratory populations from Greece, Portugal, and China. This observation suggests two possible scenarios: i) the existence of detrimental effects associated with the presence of these viruses, selecting for laboratory-adapted insects devoid of these viruses, or ii) the adaptation of existing viruses infected closely related host species after the introduction of *T. absoluta* into Spain in 2006 (Urbaneja, *et al*., 2007, Biondi *et al*., 2018). One or the other hypothesis may be virus-dependent. In favor of the first hypothesis, the laboratory colony collected in Spain and analyzed in this work, is also free of any viral infection. Nevertheless, and against this first hypothesis, multiple RNA viruses have been found simultaneously infecting laboratory insects (Shi *et al*., 2020, Hernández-Pelegrín *et al*., 2022) without an obvious impact on the fitness of the insects. Related with the second hypothesis, it is known that, although the host range of RNA viruses is relatively restricted, the same virus can be infective for closely related species (Longdon *et al*., 2011, 2015). In the case of *T. absoluta*, its invasion into new regions may have favored its exposure to previously unknown viruses.

Viral sequences obtained through metatranscriptomics not only reflect the virome of the host organism but also include viruses of associated organisms. In this study, some of the detected viruses are plant-infecting viruses and mycoviruses. The detection of plant viruses in *T. absoluta* is not surprising, as this pest feeds on the mesophyll of tomato plants and other solanaceous species (Abbes *et al*., 2016; Biondi *et al*., 2018). Notably, the widespread presence of the ToMV in nearly all evaluated samples was unexpected (Fig. 2). However, tobamoviruses are known for their remarkable environmental stability, remaining viable for extended periods outside their host plants (Rivarez *et al*., 2021). Its global distribution in our samples leads us to suspect that *T. absoluta* may serve as secondary vector for ToMV, as has been suggested for ToBRFV (Caruso *et al*., 2024), a recently emerged tobamovirus responsible for outbreaks in tomato production worldwide (Salem *et al*., 2016, 2023). Infected adults of *T. absoluta* can transmit ToBRFV to healthy tomato plants, though the virus does not circulate in the progeny of the insect (Caruso *et al*., 2024). On the other hand, the high diversity of mycoviruses identified primarily in one sample of dead larvae is noteworthy. In contrast, live larvae from the same population showed no evidence of mycoviruses, suggesting that the observed diversity may be attributed to the decomposed material (Suppl. Fig. S2). Similarly, a study of soybean thrips, *Neohydatothrips variabilis*, revealed a significant variety of viruses, including those infecting arthropods, fungi, and plants. Among these, 22 virus-like sequences were associated with mycoviruses, six of which were novel viruses (Thekke-Veetil *et al*., 2020).

In conclusion, our results provide the first insights into the composition and diversity of RNA virus present in *T. absoluta*, including ISVs as well as plant-infecting viruses and mycoviruses associated with this pest. Further investigation into virus-host interactions will be essential for validating the biological roles of these viruses and could offer valuable information for developing integrated pest management (IPM) strategies.

## Author’s contributions

**Rosa Esmeralda Becerra-García:** Writing – original draft, review & editing, Methodology, Investigation, Formal analysis, Conceptualization. **Luis Hernández-Pelegrín:** Review & editing, Methodology, Investigation, Conceptualization. **Cristina Crava:** Writing – review & editing, Supervision, Funding acquisition. **Salvador Herrero:** Writing – review & editing, Supervision, Resources, Project administration, Funding acquisition, Conceptualization.

## Acknowledgements

This study was supported by grants [PID2021-124813OB-C33], funded by MCIN/AEI/10.13039/501100011033 and by ‘ERDF A way of making Europe’, by the European Union, and by grant from the Generalitat Valenciana [CIPROM/2023/56]. REBG was supported by a Santiago Grisolía Fellowship from the Generalitat Valenciana [CIGRIS/2021/077] CMC was supported by a Ramón y Cajal grant [RYC2021-033098-I] funded by MICIU/AEI/10.13039/501100011033 and NextGenerationEU/PRTR. We want to thank Judit Arnó (IRTA, Cabrils, Spain), María Jose Pozo group (EEZ, Granada, Spain), Alberto Urbaneja Group (IVIA), and CASI cooperative (Almería, Spain) for their help with the insect collection in the field.

