## Supplementary material for "Characterization of the *Tuta absoluta* virome reveals higher viral diversity in field populations": Suppl_material.docx

Supplementary Figure S1:

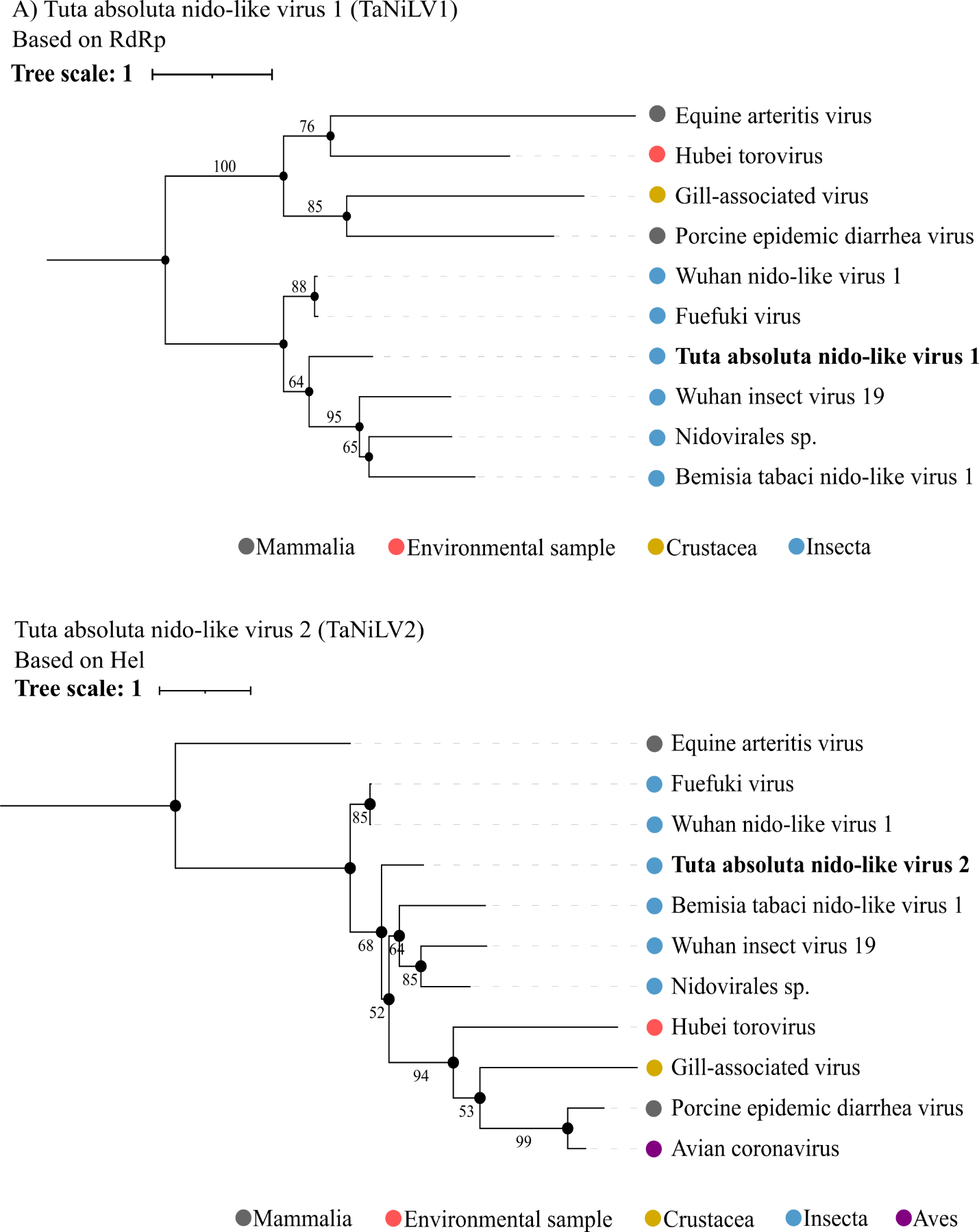

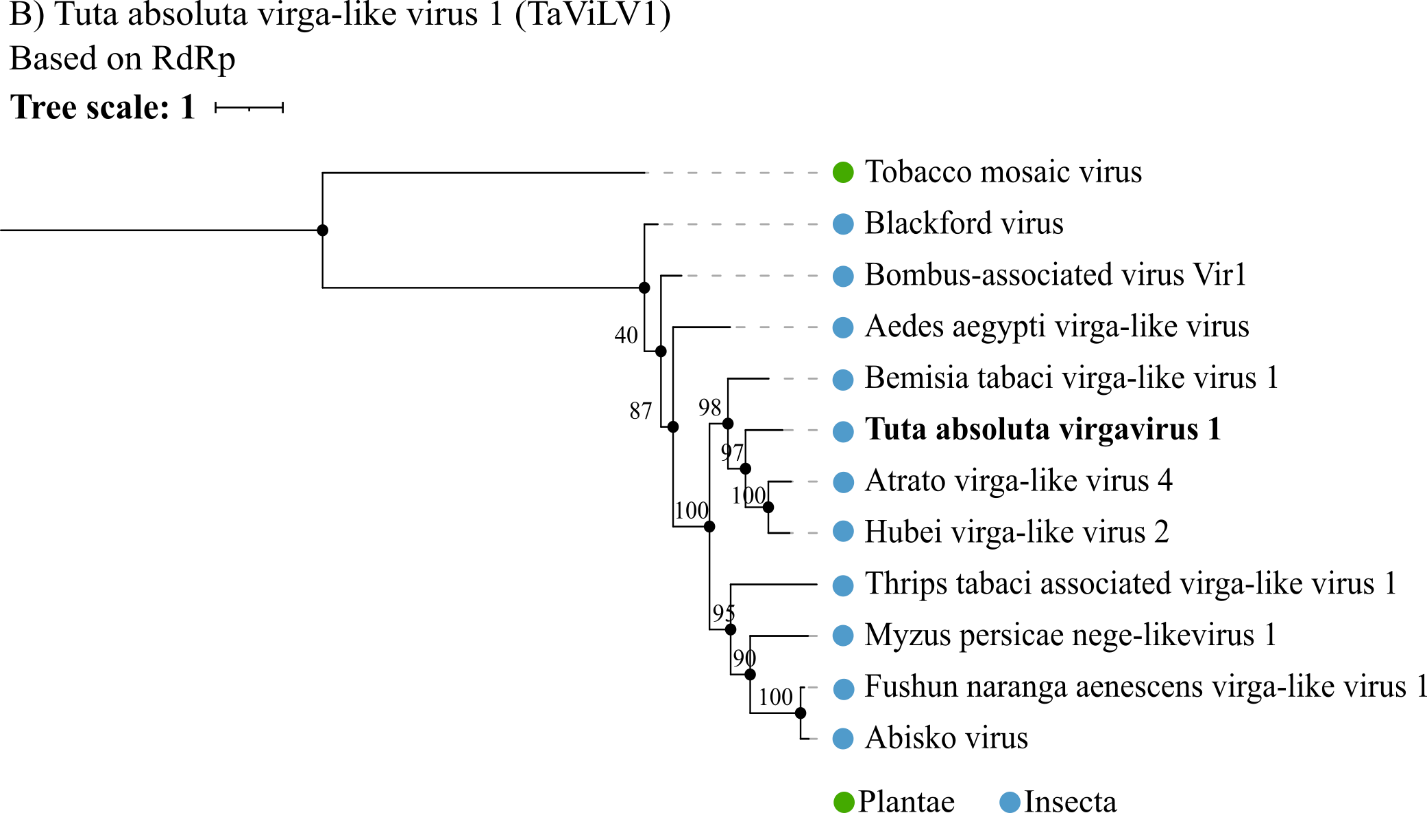

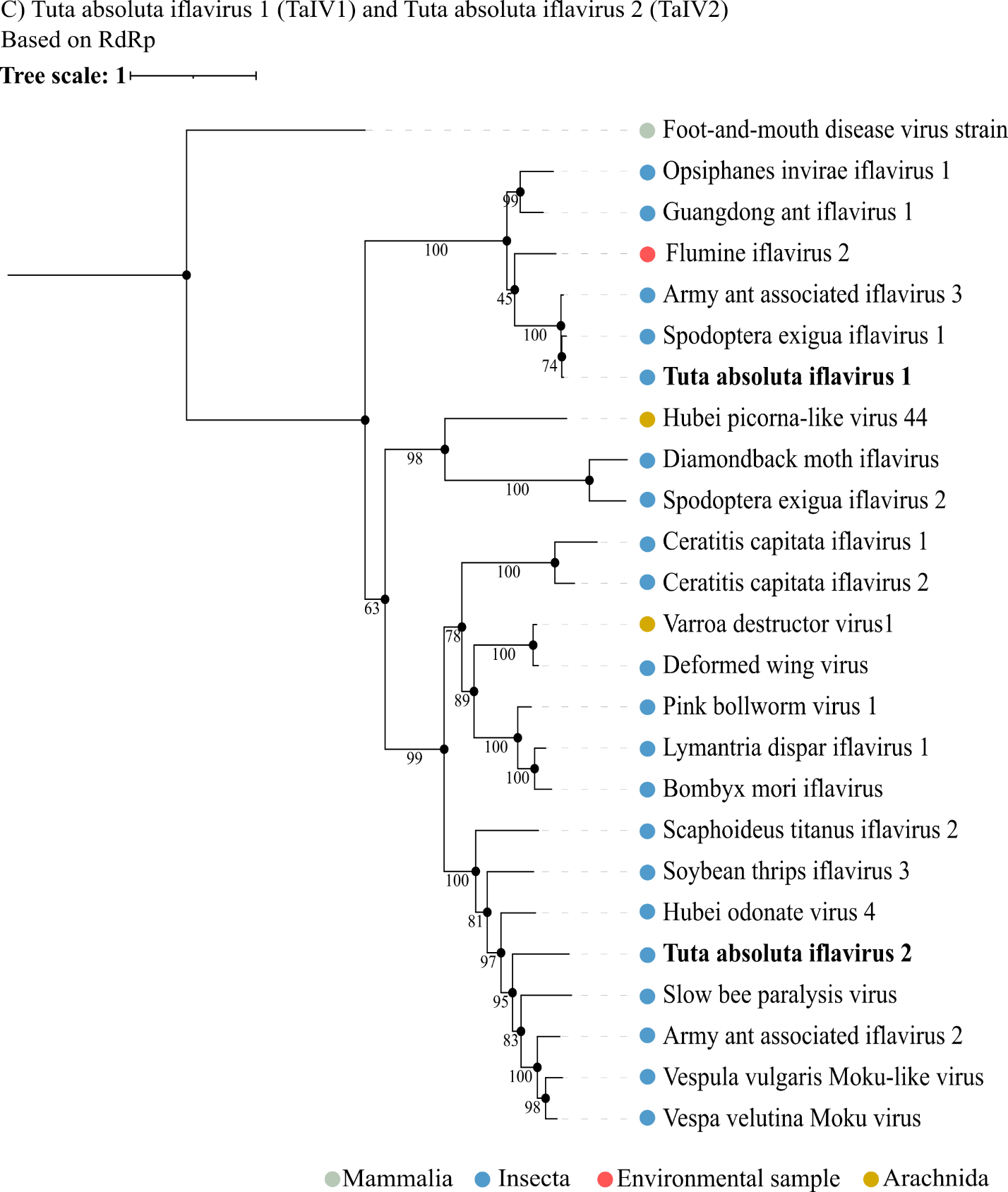

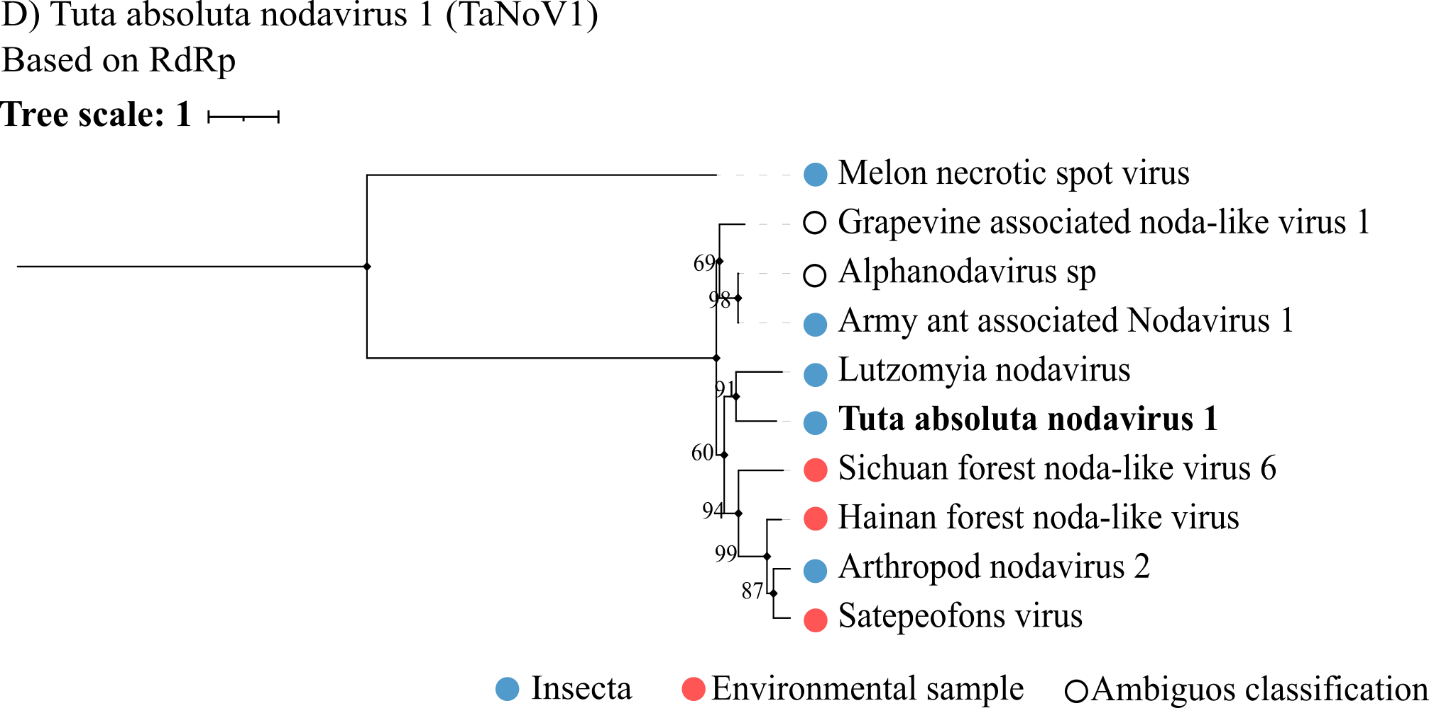

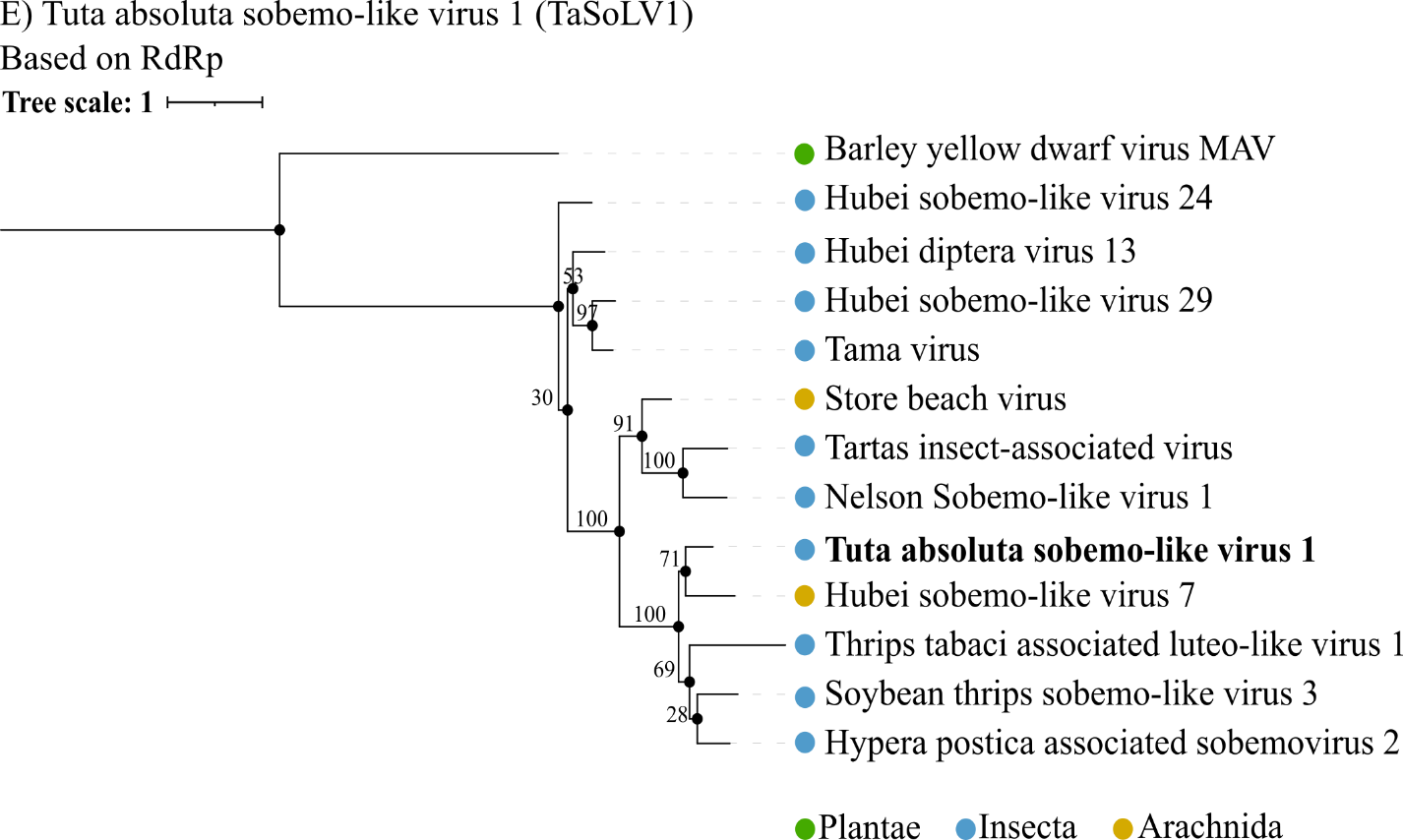

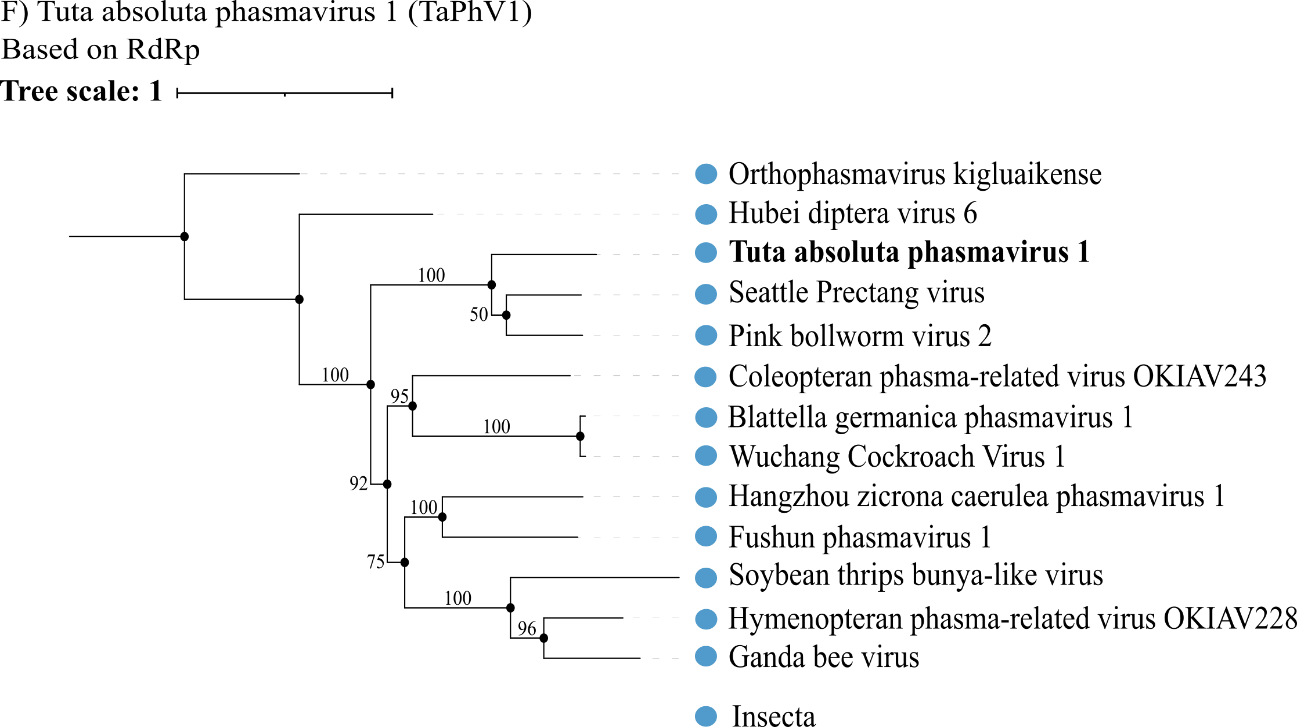

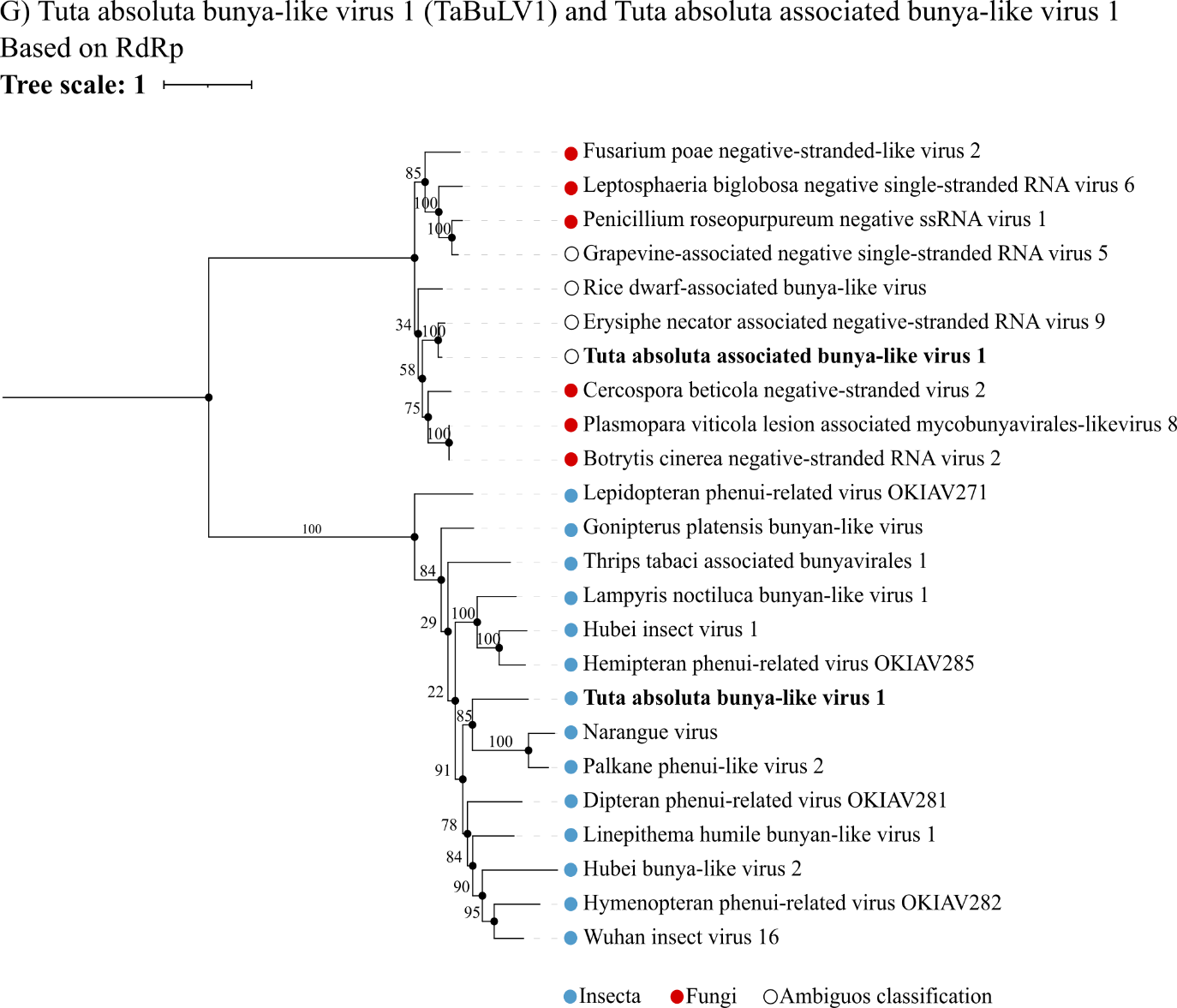

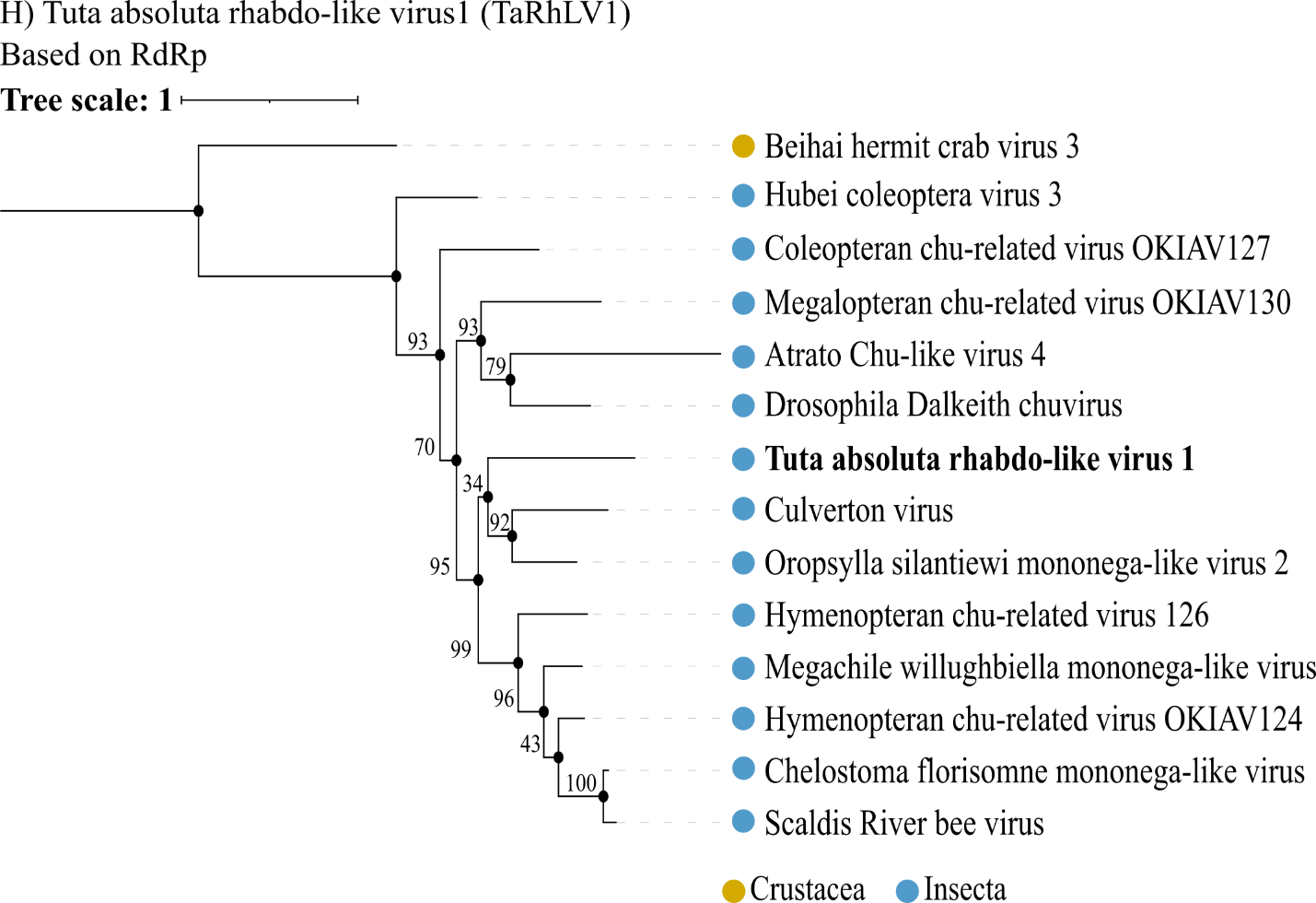

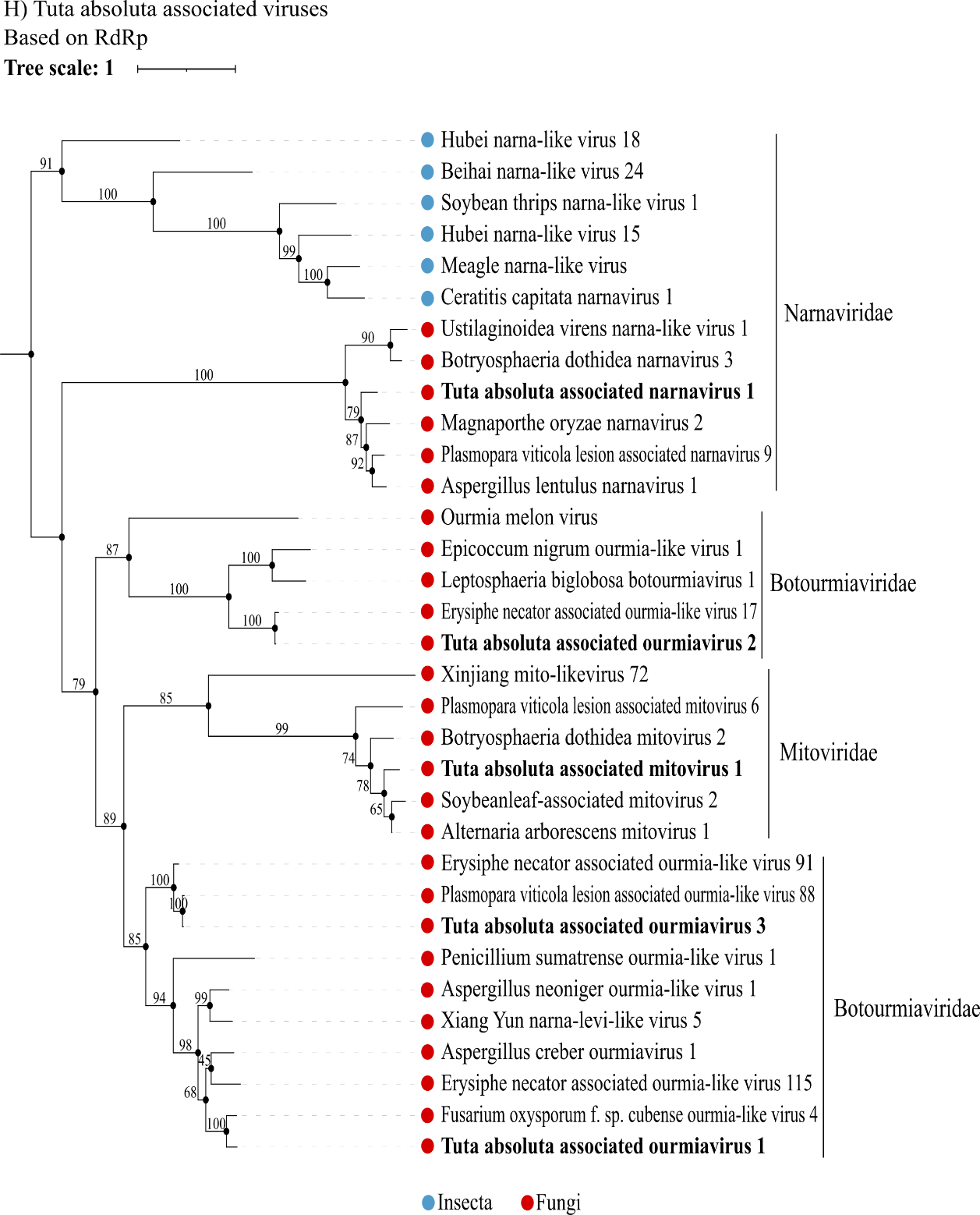

**Suppl. Fig S1. Maximum likelihood phylogenies of novel RNA viruses discovered in *T. absoluta* populations.** Trees for A) Nidovirales, B) *Virgaviridae*, C) *Iflaviridae*, D) *Nodaviridae*, E) *Solemoviridae*, F) *Phasmaviridae*, G) Bunyavirales, H) Mononegavirales and I) Novel mycoviruses associated with *T. absoluta*, were based on protein sequences of conserved viral protein domains. The protein used is indicated at the top of each tree. Branch lengths are indicated by the scale bar. The novel RNA viruses identified in this study are shown in bold font. The viral protein sequences of related viruses used in the phylogenetic analyses are listed in Suppl. Table S2.

Supplementary Figure S2:

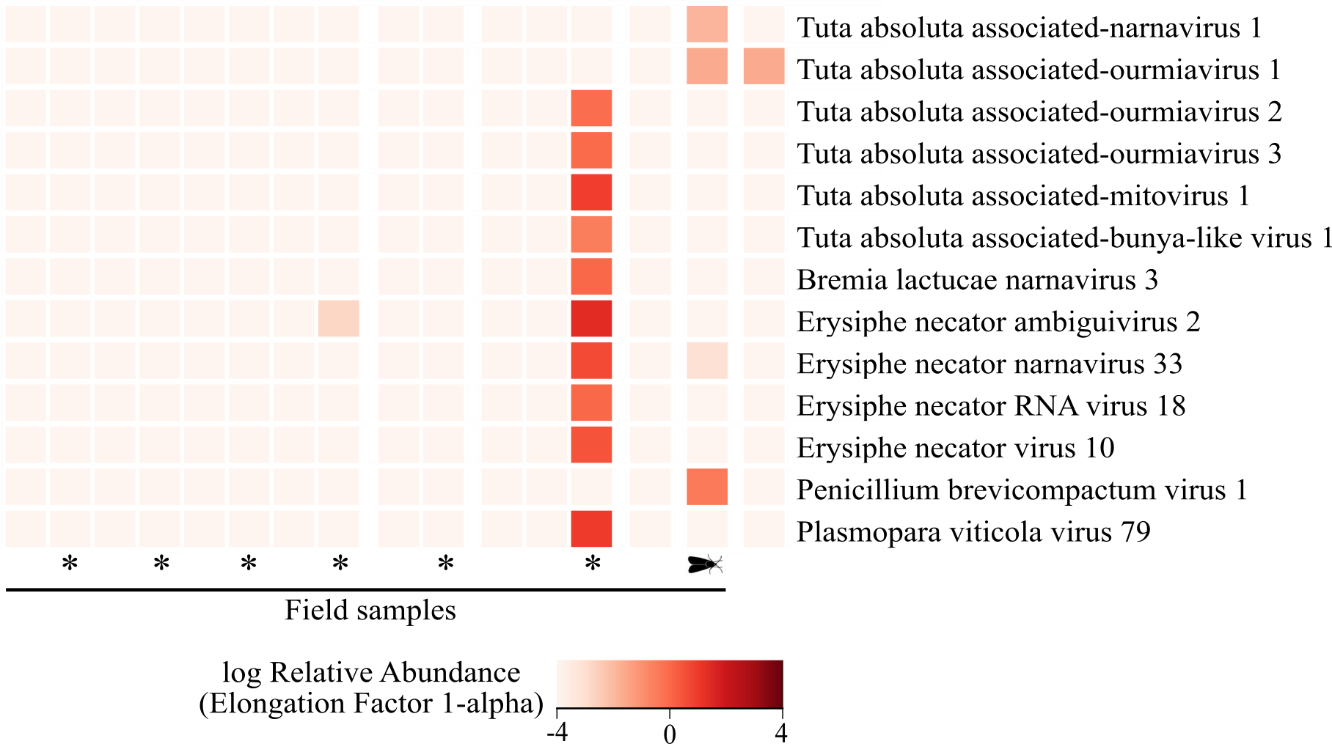

**Suppl. Fig. S2.** **Relative abundance of identified RNA mycoviruses in *Tuta absoluta***. Heatmap of the relative abundance of 13 RNA mycoviruses identified in 16 meta-transcriptomic pools corresponding to Spanish locations. The viral relative abundance was obtained by normalizing to the expression of the endogenous gene EF1α of *T. absoluta*. Columns without symbol refers to live larvae, asterisk symbol refers to dead larvae, and icon of moth refers to moth sample. The latest sample corresponds to a laboratory sample.

**Supplementary table S1.** Information of 58 *Tuta absoluta* samples used for viral discovery.

| **Bioproject** | **Origin** | **Location** | **Submitted by** | **Developmental stage** | **Biosample or SRA accession number** | **Year of collection** |
| --- | --- | --- | --- | --- | --- | --- |
| PRJNA1188764 | Field | Almeria, Spain | Universitat de València | Larvae | SAMN44848534 | 2021 |
|  |  |  |  | Larvae | SAMN44848535 |  |
|  |  |  |  | Larvae | SAMN44848536 |  |
|  |  |  |  | Larvae | SAMN44848537 |  |
|  |  |  |  | Larvae | SAMN44848538 |  |
|  |  |  |  | Larvae | SAMN44848539 |  |
|  |  |  |  | Larvae | SAMN44848540 |  |
|  |  |  |  | Larvae | SAMN44848541 |  |
|  |  | Xilxes, Spain |  | Larvae | SAMN44848542 | 2022 |
|  |  |  |  | Larvae | SAMN44848543 |  |
|  |  | Zafarraya, Spain |  | Larvae | SAMN44848544 | 2022 |
|  |  |  |  | Larvae | SAMN44848545 |  |
|  |  |  |  | Larvae | SAMN44848546 |  |
|  |  | Blanes and Quintana, Spain |  | Larvae | SAMN44848547 | 2021 |
|  |  | Multiple locations of Spain |  | Adults | SAMN44848548 | 2022 |
|  | Lab | IVIA, Spain |  | Larvae | SAMN44848549 | 2023 |
| PRJNA749726 |  | Murcia, Spain | Foundation for Research and Technology Hellas | Larvae 2^nd^ instar | [SRR15248437](https://trace.ncbi.nlm.nih.gov/Traces?run=SRR15248437) | 2019 |
|  |  |  |  |  | [SRR15248436](https://trace.ncbi.nlm.nih.gov/Traces/?view=run_browser&acc=SRR15248436&display=metadata) |  |
|  |  |  |  |  | [SRR15248445](https://trace.ncbi.nlm.nih.gov/Traces?run=SRR15248445) |  |
|  |  |  |  |  | [SRR15248444](https://trace.ncbi.nlm.nih.gov/Traces?run=SRR15248444) |  |
| PRJNA300412 |  | Cartagena, Spain | Rothamsted Research | Pool of larvae | [SRR2846714](https://trace.ncbi.nlm.nih.gov/Traces?run=SRR2846714) | 2010 |
|  |  | Portugal |  | Larvae | [SRR2913248](https://trace.ncbi.nlm.nih.gov/Traces?run=SRR2913248) | 2012 |
|  |  |  |  |  | [SRR2913250](https://trace.ncbi.nlm.nih.gov/Traces?run=SRR2913250) |  |
|  |  |  |  |  | [SRR2913254](https://trace.ncbi.nlm.nih.gov/Traces?run=SRR2913254) |  |
|  |  |  |  |  | [SRR2913258](https://trace.ncbi.nlm.nih.gov/Traces?run=SRR2913258) |  |
|  |  |  |  |  | [SRR2913261](https://trace.ncbi.nlm.nih.gov/Traces?run=SRR2913261) |  |
|  |  |  |  |  | [SRR2913263](https://trace.ncbi.nlm.nih.gov/Traces?run=SRR2913263) |  |
| PRJNA749726 |  | Greece | Foundation for Research and Technology Hellas | Larvae 2^nd^ instar | [SRR15248441](https://trace.ncbi.nlm.nih.gov/Traces?run=SRR15248441) | 2019 |
|  |  |  |  |  | [SRR15248440](https://trace.ncbi.nlm.nih.gov/Traces?run=SRR15248440) |  |
|  |  |  |  |  | [SRR15248439](https://trace.ncbi.nlm.nih.gov/Traces?run=SRR15248439) |  |
|  |  |  |  |  | [SRR15248438](https://trace.ncbi.nlm.nih.gov/Traces?run=SRR15248438) |  |
|  |  |  |  |  | [SRR15248447](https://trace.ncbi.nlm.nih.gov/Traces?run=SRR15248447) |  |
|  |  |  |  |  | [SRR15248446](https://trace.ncbi.nlm.nih.gov/Traces?run=SRR15248446) |  |
|  |  |  |  |  | [SRR15248443](https://trace.ncbi.nlm.nih.gov/Traces?run=SRR15248443) |  |
|  |  |  |  |  | [SRR15248442](https://trace.ncbi.nlm.nih.gov/Traces?run=SRR15248442) |  |
| PRJNA291932 |  | Brazil | Pontifícia Universidade Católica do Paraná | Larvae 2^nd^ instar | [SRR2147319](https://trace.ncbi.nlm.nih.gov/Traces/?view=run_browser&acc=SRR2147319&display=metadata) | 2011 |
|  |  |  |  | Larvae 3^rd^ instar | [SRR2147320](https://trace.ncbi.nlm.nih.gov/Traces/?view=run_browser&acc=SRR2147320&display=metadata) |  |
|  |  |  |  | Larvae 4^th^ instar | [SRR2147321](https://trace.ncbi.nlm.nih.gov/Traces/?view=run_browser&acc=SRR2147321&display=metadata) |  |
|  |  |  |  | Adults | [SRR2147322](https://trace.ncbi.nlm.nih.gov/Traces/?view=run_browser&acc=SRR2147322&display=metadata) |  |
|  |  |  |  | Eggs | [SRR2147323](https://trace.ncbi.nlm.nih.gov/Traces/?view=run_browser&acc=SRR2147323&display=metadata) |  |
| PRJNA869533 |  | China | Guizhou University | Larvae 3^rd^ instar | [SRR21101434](https://trace.ncbi.nlm.nih.gov/Traces?run=SRR21101434) | Undetermined |
|  |  |  |  |  | [SRR21101435](https://trace.ncbi.nlm.nih.gov/Traces?run=SRR21101435) |  |
|  |  |  |  |  | [SRR21101436](https://trace.ncbi.nlm.nih.gov/Traces?run=SRR21101436) |  |
|  |  |  |  |  | [SRR21101437](https://trace.ncbi.nlm.nih.gov/Traces?run=SRR21101437) |  |
|  |  |  |  |  | [SRR21101438](https://trace.ncbi.nlm.nih.gov/Traces?run=SRR21101438) |  |
|  |  |  |  |  | [SRR21101439](https://trace.ncbi.nlm.nih.gov/Traces?run=SRR21101439) |  |
|  |  |  |  |  | [SRR21101440](https://trace.ncbi.nlm.nih.gov/Traces?run=SRR21101440) |  |
|  |  |  |  |  | [SRR21101441](https://trace.ncbi.nlm.nih.gov/Traces?run=SRR21101441) |  |
|  |  |  |  |  | [SRR21101442](https://trace.ncbi.nlm.nih.gov/Traces?run=SRR21101442) |  |
|  |  |  |  |  | [SRR21101443](https://trace.ncbi.nlm.nih.gov/Traces?run=SRR21101443) |  |
|  |  |  |  |  | [SRR21101444](https://trace.ncbi.nlm.nih.gov/Traces?run=SRR21101444) |  |
|  |  |  |  |  | [SRR21101445](https://trace.ncbi.nlm.nih.gov/Traces?run=SRR21101445) |  |
| PRJNA926790 |  | Yunnan, China | China Agricultural University (Yunnan Academy of Agricultural Sciences) | Pupa | [SRR23345409](https://trace.ncbi.nlm.nih.gov/Traces/?view=run_browser&acc=SRR23345409&display=metadata) | 2021 |
|  |  |  |  | Larvae 4^th^ instar | [SRR23345410](https://trace.ncbi.nlm.nih.gov/Traces/?view=run_browser&acc=SRR23345410&display=metadata) |  |
|  |  |  |  | Larvae 3^rd^ instar | [SRR23345411](https://trace.ncbi.nlm.nih.gov/Traces/?view=run_browser&acc=SRR23345411&display=metadata) |  |
|  |  |  |  | Larvae 2^nd^ instar | [SRR23345412](https://trace.ncbi.nlm.nih.gov/Traces/?view=run_browser&acc=SRR23345412&display=metadata) |  |
|  |  |  |  | Larvae 1^st^ instar | [SRR23345413](https://trace.ncbi.nlm.nih.gov/Traces/?view=run_browser&acc=SRR23345413&display=metadata) |  |
| PRJNA678630 |  | China | Guiyang University | Undetermined | [SRR13065833](https://trace.ncbi.nlm.nih.gov/Traces/?view=run_browser&acc=SRR13065833&display=metadata) | 2020 |

Samples are presented in the same order as in Fig. 2, which includes the shadow samples.

**Supplementary Table S2**. List of viral proteins included in the phylogenetic analysis.

| **Viral name** | **Protein accession number** | **Viral name** | **Protein accession number** |
| --- | --- | --- | --- |
| (A) Nidovirales | | (C) *Iflaviridae* | |
| Tuta absoluta nido-like virus 1 | PQ655384* | Tuta absoluta iflavirus 1 | PQ655387* |
| Tuta absoluta nido-like virus 2 | PQ655385* | Tuta absoluta iflavirus 2 | PQ655388* |
| Equine arteritis virus | QID92065.1 | Foot-and-mouth disease virus (strain O1) | AAF09193.1 |
| Hubei torovirus | QYF49656.1 | Opsiphanes invirae iflavirus 1 | YP_009167346.1 |
| Gill-associated virus | YP_001661452.1 | Guangdong ant iflavirus 1 | WXH83723.1 |
| Porcine epidemic diarrhea virus | UKF18845.1 | Flumine iflavirus 2 | UQB76023.1 |
| Wuhan nido-like virus 1 | YP_009345058.1 | Army ant associated iflavirus 3 | WAX26121.1 |
| Fuefuki virus | AWA82245.1 | Spodoptera exigua iflavirus 1 | YP_004935363.1 |
| Wuhan insect virus 19 | YP_009342322.1 | Hubei picorna-like virus 44 | WAK72349.1 |
| Nidovirales sp. | URA30378.1 | Diamondback moth iflavirus | YP_009361829.1 |
| Bemisia tabaci nido-like virus 1 | QWC36473.1 | Spodoptera exigua iflavirus 2 | YP_009010984.1 |
| Avian coronavirus | WBW48761.1 | Ceratitis capitata iflavirus 1 | JAC04636.1 |
| (B) *Virgaviridae* | | Ceratitis capitata iflavirus 2 | UOI84715.1 |
| Tuta absoluta virga-like virus 1 | PQ655386* | Varroa destructor virus 1 | WJN61826.1 |
| Tobacco mosaic virus | ABN79256.1 | Deformed wing virus | WDY79011.1 |
| Blackford virus | AMO03220.1 | Pink bollworm virus 1 | QID77674.1 |
| Bombus-associated virus Vir1 | QAY29261.1 | Lymantria dispar iflavirus 1 | YP_009047245.1 |
| Aedes aegypti virga-like virus | BBN20999.1 | Bombyx mori iflavirus | YP_009162630.1 |
| Bemisia tabaci virga-like virus 1 | QNJ34552.1 | Scaphoideus titanus iflavirus 2 | QIJ56911.1 |
| Atrato Virga-like virus 4 | QHA33746.1 | Soybean thrips iflavirus 3 | QQN90112.1 |
| Hubei virga-like virus 2 | UYE93717.1 | Hubei odonate virus 4 | YP_009337760.1 |
| Thrips tabaci associated virga-like virus 1 | QNM37812.1 | Slow bee paralysis virus | YP_003622540.1 |
| Myzus persicae nege-like virus 1 | UTQ79656.1 | Army ant associated iflavirus 2 | WAX26119.1 |
| Fushun naranga aenescens virga-like virus 1 | UHM27606.1 | Vespula vulgaris Moku-like virus | QZZ63303.1 |
| Abisko virus | YP_009408586.1 | Vespa velutina Moku virus | ATY36108.1 |

| **Viral name** | **Protein accession number** | **Viral name** | **Protein accession number** |
| --- | --- | --- | --- |
| (D) *Nodaviridae* | | (G) Bunya-like viruses | |
| Tuta absoluta nodavirus 1 | PQ655389* | Tuta absoluta bunya-like virus 1 | PQ655396* |
| Melon necrotic spot virus | APG76350.1 | Tuta absoluta associated-bunya-like virus 1 | PQ655398* |
| Grapevine-associated noda-like virus 1 | QXN75416.1 | Fusarium poae negative-stranded-like virus 2 | QQO58802.1 |
| Alphanodavirus sp. | XCO48224.1 | Leptosphaeria biglobosa negative single-stranded RNA virus 6 | UYL94507.1 |
| Army ant associated Nodavirus 1 | WAX26160.1 | Penicillium roseopurpureum negative ssRNA virus 1 | YP_010840311.1 |
| Lutzomyia nodavirus | AKP18615.1 | Grapevine-associated negative single-stranded RNA virus 5 | QXN75415.1 |
| Sichuan forest noda-like virus 6 | QYF49974.1 | Rice dwarf-associated bunya-like virus | UTJ93941.1 |
| Hainan forest noda-like virus | QYF49887.1 | Erysiphe necator associated negative-stranded RNA virus 9 | QJW70358.1 |
| Arthropod nodavirus 2 | WPR16578.1 | Cercospora beticola negative-stranded virus 2 | UVB78666.1 |
| Satepeofons virus | WNT71152.1 | Plasmopara viticola lesion associated mycobunyavirales-like virus 8 | YP_010840346.1 |
| (E) *Solemoviridae* | | Botrytis cinerea negative-stranded RNA virus 2 | QJT73695.1 |
| Tuta absoluta sobemo-like virus 1 | PQ655391* | Lepidopteran phenui-related virus OKIAV271 | QMP82187.1 |
| Barley yellow dwarf virus MAV | NP_620064.1 | Gonipterus platensis bunyan-like virus | QWX94187.1 |
| Hubei sobemo-like virus 24 | YP_009330073.1 | Thrips tabaci associated bunyavirales 1 | QNS31055.1 |
| Hubei diptera virus 13 | YP_009337911.1 | Lampyris noctiluca bunyan-like virus 1 | QBP37023.1 |
| Hubei sobemo-like virus 29 | YP_009330084.1 | Hubei insect virus 1 | APG79218.1 |
| Tama virus | AWA82270.1 | Hemipteran phenui-related virus OKIAV285 | QMP82212.1 |
| Store beach virus | AYP67537.1 | Narangue virus | YP_010839995.1 |
| Tartas insect-associated virus | UNZ11822.1 | Palkane phenui-like virus 2 | UYE93926.1 |
| Nelson Sobemo-like virus 1 | QZZ63405.1 | Dipteran phenui-related virus OKIAV281 | QMP82130.1 |
| Hubei sobemo-like virus 7 | YP_009330005.1 | Linepithema humile bunyan-like virus 1 | AXA52548.1 |
| Thrips tabaci associated luteo-like virus 1 | QNM37821.1 | Hubei bunya-like virus 2 | APG79269.1 |
| Soybean thrips sobemo-like virus 3 | QPZ88397.1 | Hymenopteran phenui-related virus OKIAV282 | QMP82348.1 |
| Hypera postica associated sobemovirus 2 | QUS52862.1 | Wuhan insect virus 16 | APG79216.1 |
| (F) *Phasmaviridae* | | | |
| Tuta absoluta phasmavirus 1 | PQ655392* | Wuchang Cockroach Virus 1 | YP_009304995.1 |
| Orthophasmavirus kigluaikense | YP_009362029.1 | Hangzhou zicrona caerulea phasmavirus 1 | UHK03218.1 |
| Hubei diptera virus 6 | UDL14021.1 | Fushun phasmavirus 1 | YP_010840784.1 |
| Seattle Prectang virus | YP_009666959.1 | Soybean thrips bunya-like virus 3 | QQX28936.1 |
| Pink bollworm virus 2 | QID77675.1 | Hymenopteran phasma-related virus OKIAV228 | YP_010840616.1 |
| Coleopteran phasma-related virus OKIAV243 | QMP82227.1 | Ganda bee virus | YP_009666981.1 |
| Blattella germanica phasmavirus 1 | DBA56461.1 |  |  |

| **Viral name** | **Protein accession number** | **Viral name** | **Protein accession number** |
| --- | --- | --- | --- |
| (H) Rhabdo-like viruses | | (I) ISVs and mycoviruses | |
| Tuta absoluta rhabo-like virus 1 | PQ655395* | Meagle narna-like virus | QIJ70070.1 |
| Beihai hermit crab virus 3 | YP_009333157.1 | Ceratitis capitata narnavirus 1 | UOI84716.1 |
| Hubei coleoptera virus 3 | YP_009336866.1 | Ustilaginoidea virens narna-like virus 1 | UVX28909.1 |
| Coleopteran chu-related virus OKIAV127 | QMP82184.1 | Botryosphaeria dothidea narnavirus 3 | QQD86177.1 |
| Megalopteran chu-related virus OKIAV130 | QPL15366.1 | Magnaporthe oryzae narnavirus 2 | BCH36659.1 |
| Atrato Chu-like virus 4 | QHA33906.1 | Plasmopara viticola lesion associated narnavirus 9 | QIR30288.1 |
| Drosophila Dalkeith chuvirus | WPV74294.1 | Aspergillus lentulus narnavirus 1 | BCH36643.1 |
| Culverton virus | YP_010798359.1 | Ourmia melon virus | ACF16360.1 |
| Oropsylla silantiewi mononega-like virus 2 | YP_010802321.1 | Epicoccum nigrum ourmia-like virus 1 | YP_010798247.1 |
| Hymenopteran chu-related virus 126 | YP_010798596.1 | Leptosphaeria biglobosa botourmiavirus 1 | UYL94487.1 |
| Megachile willughbiella mononega-like virus | DAZ89735.1 | Erysiphe necator associated ourmia-like virus 17 | QKI79847.1 |
| Hymenopteran chu-related virus OKIAV124 | QPL15325.1 | Xinjiang mito-like virus 72 | UPW42227.1 |
| Chelostoma florisomne mononega-like virus | DAZ89733.1 | Plasmopara viticola lesion associated mitovirus 6 | QIR30230.1 |
| Scaldis River bee virus | UDL14020.1 | Botryosphaeria dothidea mitovirus 2 | QUP79244.1 |
| (I) ISVs and mycoviruses | | Soybean leaf-associated mitovirus 2 | ALM62242.1 |
| Tuta absoluta associated-mitovirus 1 | PQ655399* | Alternaria arborescens mitovirus 1 | QJT93894.1 |
| Tuta absoluta associated-narnavirus 1 | PQ655400* | Erysiphe necator associated ourmia-like virus 91 | QKI79921.1 |
| Tuta absoluta associated-ourmiavirus 1 | PQ655401* | Plasmopara viticola lesion associated ourmia-like virus 88 | YP_010800281.1 |
| Tuta absoluta associated-ourmiavirus 2 | PQ655402* | Penicillium sumatrense ourmia-like virus 1 | YP_010798244.1 |
| Tuta absoluta associated-ourmiavirus 3 | PQ655403* | Aspergillus neoniger ourmia-like virus 1 | YP_010798179.1 |
| Hubei narna-like virus 18 | APG77102.1 | XiangYun narna-levi-like virus 5 | UUG74239.1 |
| Beihai narna-like virus 24 | YP_009333245.1 | Aspergillus creber ourmiavirus 1 | BDB16252.1 |
| Soybean thrips narna-like virus 1 | QQP18719.1 | Erysiphe necator associated ourmia-like virus 115 | QKI79944.1 |
| Hubei narna-like virus 15 | YP_009337783.1 | Fusarium oxysporum f. sp. cubense ourmia-like virus 4 | WNK16431.1 |

*Nucleotide accession numbers were provided when protein accession numbers were not available, and nucleotide sequences were translated into protein using the standard genetic code.
